# Tissue-specific cancer paradox and mutation rates

**DOI:** 10.1101/012658

**Authors:** Valentina Agoni

## Abstract

Some genes have ubiquitous expression patterns in an individual’s cells, thus a paradox exists whereby mutations in such genes are only strongly associated with cancers of specific tissues. As these genes are ubiquitously expressed in the body’s cells and thereby have functions in -potentially- all tissue types, then surely their resulting defects would manifest as cancers in all tissues?

We hypothesize that the different susceptibility to mutations could explain the ‘tissue-specific’ paradox for cancer.

---

Tumors derive from mutations in proto-oncogenes and oncosuppressors. Proto-oncogenes regulate cell-cycle progression. Mutations of these genes cause an abnormal proliferation that leads to cancer. On the other hand non-functioning oncosuppressors could result in DNA damage accumulation and possible neoplastic transformation.

There are certain genes that have ubiquitous expression patterns in an individual’s cells; yet, a paradox exists whereby mutations (germline or somatic) in such genes are only strongly associated with cancers of specific tissues. As these genes are ubiquitously expressed in the body’s cells and thereby have functions in -potentially- all tissue types, then surely their resulting defects would manifest as cancers in all tissues? (Blighe, 2014).

Contemporary, in the last years the correlation with tissue-specific mutations emerged for different conditions such as genetic diseases and cancers. Recurrent tissue-specific mtDNA mutations had been found in humans (Samuels et al., 2013). Muscle-specific mutations accumulate with aging in critical human mtDNA control sites for replication (Wang et al., 2001). The recently discovered aging-dependent large accumulation of point mutations in the human fibroblast mtDNA control region raised the question of their occurrence in postmitotic tissues. In addition, mutations and tissue-specific alterations of RPGR transcripts had been described in the X-chromosome linked retinitis pigmentosa (Schmid et al., 2010) and idiopathic atrial fibrillation (Gollob et al., 2006). Hu (2009) and Blighe (2014) explore the ‘tissue-specific’ paradox for cancer genes. Sieber et al. (2005) previously analyzed the role of tissue, cell and stage specificity of (epi)mutations in cancers. They hypothesize that: (i) most (epi)mutations in cancers are specific to particular tumours or occur at specific stages of development, cell differentiation or tumorigenesis. (ii) simple molecular mechanisms, such as tissue restricted gene expression, seem to explain these associations only in rare cases. (iii) instead, the specificity of (epi)mutations is probably due to the selection of a restricted spectrum of genetic changes by the cellular environment. (iv) in some cases, the resulting functional defects might be constrained to be neither too strong nor too weak for tumour growth to occur; that is, they lie within a ‘window’ that is permissive for tumorigenesis.

**Figure 1.**
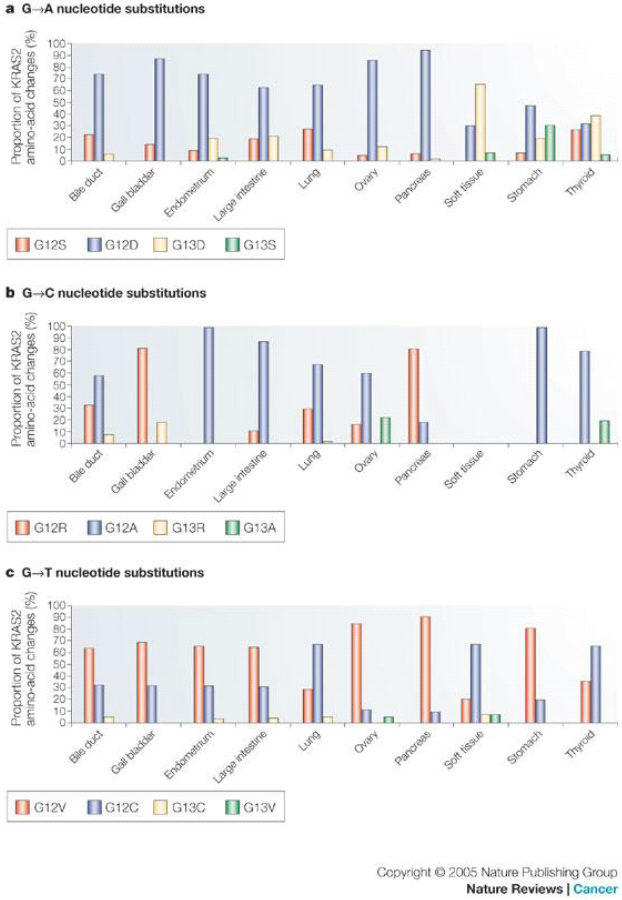
Distribution of activating *KRAS2* mutations in different tumour types. Proportions of KRAS2 amino-acid changes in different cancer types resulting from (**a**) G→A nucleotide substitutions, (**b**) G→C nucleotide substitutions, and (**c**) G→T nucleotide substitutions. Even after correcting for mutagenic bias in different tissues at the nucleotide level, activating *KRAS2* mutations still differ considerably between cancer types as regards the position (codon 12 or 13) and type of amino-acid substitution (P<0.0001 each for G→A, G→T and G→C nucleotide substitutions; chi-square test and bootstrap simulation data). Data from COSMIC database (Sieber et al., 2005).

It is known that the amino acids coding table evolved to minimize the impact of deleterious mutations. The observation of the third nucleotide of codons lead us to hypothesize that G↔A and C↔T mutations are the more probable single nucleotide substitutions (Agoni, 2013).

This is supported by the statistics for missense mutations derived from the Human Gene Mutation Database (HGMD) (Table 1).

**Table 1.**
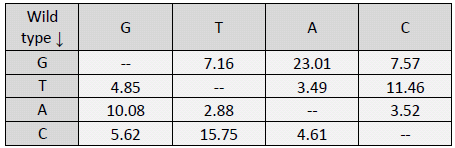
Statistics for missense mutations in percentage. Modified from The Human Gene Mutation Database at the Institute of Medical Genetics in Cardiff http://www.hgmd.cf.ac.uk/ac/hoho1.php

We do not know the rate of spontaneous mutations vs errors induced by cancerogenic agents. Nevertheless, the different percentages of mutations could explain the ‘tissue-specific’ paradox for cancer. In fact, the gene sequences in diverse tissues and diverse subjects could have a different sensibility to mutations.

In other words if the genes expressed in a particular tissue are GC-richest respect to another, they are more prone to mutations, thus the tissues are differentially exposed to neoplastic transformations too.

Moreover, why for example *brca1* mutates and gives tumors preferentially in female breast?

As elucidated by Hu (2009) BRCA1 recovers many functions related to estrogen/progesterone signaling pathways in addition to its role in genome stability.

We hypothesize that -assuming that genes not protected by the chromatin apparatus are more susceptible to cancerigenic mutations- the nucleotide composition of genes expressed in this tissue make *brca1* to be more exposed to mutagens in breast respect to other tissues.

Suppose that we are considering a mutagen that gives G to A mutations, if the genes expressed in the brain have an higher G-content respect to the ones expressed in the breast, consequently a G nucleotide belonging to *brca1* in the breast have an higher probability to mutate into A.

Effectively, despite very difficult to verify due to very poor data about tissue-specific transcriptome analysis and GC-content, if we focus our attention on the highly expressed genes in different tissues, we can notice (Table 2) that the tissues more exposed to cancer present in average a lower GC-content (compatibly with the great variability and with the function and lenght of the genes). In fact the most common types of cancer include breast and prostate cancers, leukemia and lymphoma (http://www.cancer.gov/cancertopics/types/commoncancers). Moreover, Stephens (2012) paper goes in accordance with our hypothesis.

**Table 2.** GC-content in the highly expressed genes. Data from Tissue-specific Gene Expression and Regulation (TiGER)

| breast |  | lung |  |
| --- | --- | --- | --- |
| gene name | GC-content (%) | gene name | GC-content (%) |
| DCD1 | 49.37 | SFTB | 53.64 |
| SCGB2 | 41.78 | SFTC | 59.96 |
| ANKRD | 38.9 | KCN | 45.97 |
| mean | 43.35 | T | 56.96 |
| pancreas |  | TITF1 | 57.65 |
| UCN3 | 61.27 | mean | 54.836 |
| IPF1 | 59.04 | colon |  |
| INS | 63.3 | KRT | 56.5 |
| REG1B | 47.66 | EVX1 | 68.35 |
| mean | 57.8175 | CDX1 | 62.75 |
| bone marrow |  | APOBEC1 | 44.35 |
| IL23R | 39.6 | GPR | 64 |
| FLJ | 37 | mean | 59.19 |
| CLC | 44.38 | prostate |  |
| OR51 | 44.44 | UPK3A | 59.56 |
| FLT3 | 43.06 | SEMG1 | 41.31 |
| FAM | 57.42 | PRAC | 46.4 |
| MS4 | 40.66 | MSMB | 43.07 |
| mean | 43.79428571 | KLK2 | 50.82 |
| ovary |  | mean | 48.232 |
| ASP | 61.5 | muscle |  |
| TTC | 59.6 | HTN | 34.78 |
| TOR2A | 60.2 | MYOD | 64.78 |
| SOX17 | 58.13 | MYLK2 | 59.7 |
| mean | 59.8575 | MUC7 | 40.17 |
|  |  | mean | 49.8575 |

At present, we know little about the mutational processes responsible for the generation of somatic mutations in breast and other cancers. In the 100 breast cancers analysed by Stephens and colleagues (2012), there was substantial variation in the total numbers of base substitutions and indels between individual cases. There was also considerable diversity of mutational pattern, ranging from cases in which C•G→T•A transitions predominated to cases in which all transitions and transversions made equal contributions. Taken together, they suggest that multiple distinct mutational processes are operative. However, for most of these processes, the underlying mechanism is unknown. To characterize this process further, they examined the sequence context in which the mutations occurred (Stephens et al., 2012).

The different sensibility to mutations could give reason also for dominate-recessive (gain-loss of function) phenotypes or it could plays a role in epigenetic. Moreover some tissue are more exposed to reactive oxygen species (ROS) mutations and we know that the effects of ROS inside cells are still matter of debate. Finally, the knowledge of missense mutations rates could help in preventing drug-resistance in bacteria, for example if we are able to predict the mutation that determines the resistance against a drug we could create a second drug so that the mutation that confers resistance to this second molecule restores the beginning situation.

